# Priming in a permissive type I-C CRISPR-Cas system reveals distinct dynamics of spacer acquisition and loss

**DOI:** 10.1101/137067

**Authors:** Chitong Rao, Denny Chin, Alexander W. Ensminger

## Abstract

CRISPR-Cas is a bacterial and archaeal adaptive immune system that uses short, invader-derived sequences termed spacers to target invasive nucleic acids. Upon recognition of previously encountered invaders, the system can stimulate secondary spacer acquisitions, a process known as primed adaptation. Previous studies of primed adaptation have been complicated by intrinsically high interference efficiency of most systems against *bona fide* targets. As such, most primed adaptation to date has been studied within the context of imperfect sequence complementarity between spacers and targets. Here, we take advantage of a native type I-C CRISPR-Cas system in *Legionella pneumophila* that displays robust primed adaptation even within the context of a perfectly matched target. Using next-generation sequencing to survey acquired spacers, we observe strand bias and positional preference that are consistent with a 3′ to 5′ translocation of the adaptation machinery. We show that spacer acquisition happens in a wide range of frequencies across the plasmid, including a remarkable hotspot that predominates irrespective of the priming strand. We systematically characterize protospacer sequence constraints in both adaptation and interference and reveal extensive flexibilities regarding the protospacer adjacent motif in both processes. Lastly, in a strain with a genetically truncated CRISPR array, we observe greatly increased interference efficiency coupled with a dramatic shift away from spacer acquisition towards spacer loss. Based on these observations, we propose that the *Legionella* type I-C system represents a powerful model to study primed adaptation and the interplay between CRISPR interference and adaptation.

## Introduction

Bacteria and archaea constantly interact with mobile genetic elements including bacteriophages, plasmids, transposons and other conjugative elements (Burrus and Waldor 2004, Frost, Leplae et al. 2005). With their genomes greatly shaped by these mobile elements, these microbes can benefit from acquisition of foreign DNA, but also suffer detrimental effects from “selfish” elements such as lytic bacteriophages. To combat deleterious horizontal gene transfer, bacteria and archaea harbor multiple resistance mechanisms exemplified by CRISPR-Cas (clustered regularly interspaced short palindromic repeats and CRISPR-associated genes) systems (Labrie, Samson et al. 2010). To date, CRISPR-Cas systems have been identified in about half of genome-sequenced bacteria and archaea and include multiple types that each use distinct protein compositions to function as adaptive immunity against invasive nucleic acids (Horvath and Barrangou 2010, Marraffini and Sontheimer 2010, Makarova, Haft et al. 2011, Makarova, Wolf et al. 2015). A CRISPR array consists of distinct short spacers separated by repeat sequences and is transcribed as a non-coding RNA that undergoes further processing by Cas proteins to form individual repeat-spacer units (crRNAs). These crRNAs are loaded as guide sequences into Cas-crRNA interference complexes that bind to targeted nucleic acids (termed protospacers) with an appropriate protospacer adjacent motif (PAM) and mediate their destruction by recruiting the Cas3 nuclease (in type I systems) or by its intrinsic nuclease activity (in other types), a process known as interference (van der Oost, Westra et al. 2014, Marraffini 2015). One key feature of CRISPR-Cas immunity is its ability to adapt to new threats through the acquisition of new spacers derived from non-productive bacteriophage infection or other encounters with foreign DNA. Spacer acquisition can be either “naïve” (where the invader has not been previously cataloged in the array) or “primed” (where upon recognition of invaders previously targeted by CRISPR-Cas, secondary spacers are acquired in order to enhance protection). Compared with naïve adaptation, primed adaptation is much more efficient and reliant on recruitment of the interference machinery to a pre-existing target (Heler, Marraffini et al. 2014, Vorontsova, Datsenko et al. 2015, Staals, Jackson et al. 2016). When coupled, CRISPR interference and adaptation can effectively protect against evolving invasive elements (Andersson and Banfield 2008, Paez-Espino, Morovic et al. 2013, Paez-Espino, Sharon et al. 2015, van Houte, Ekroth et al. 2016).

Our understanding of primed spacer acquisition is based upon the studies of type I-E, type I-F and type I-B systems in the presence of targeted DNA such as plasmids and bacteriophages (Fineran and Charpentier 2012, Heler, Marraffini et al. 2014, Amitai and Sorek 2016, Sternberg, Richter et al. 2016, Jackson, McKenzie et al. 2017). During CRISPR adaptation, the conserved proteins Cas1 and Cas2 form a protein complex that plays a key role in pre-spacer capture and insertion into the CRISPR array (Nunez, Kranzusch et al. 2014, Nunez, Harrington et al. 2015, Nunez, Lee et al. 2015, Wang, Li et al. 2015). Regarding the generation of pre-spacers from invasive DNA, characterization of acquired spacers in a priming condition revealed non-conserved patterns in different type I CRISPR-Cas systems. Primed spacer acquisition in the *E. coli* type I-E system showed a clear preference (>90%) from the primed (untargeted) strand and no obvious positional gradient on the plasmid or bacteriophage (Datsenko, Pougach et al. 2012, Savitskaya, Semenova et al. 2013, Fineran, Gerritzen et al. 2014). In the *Haloarcula hispanica* type I-B system, ∼70% of new spacers were derived from the primed strand and a moderate preference was seen for the priming-proximal region (Li, Wang et al. 2014). In the *Pectobacterium atrosepticum* type I-F system, ∼65% of spacers were acquired from the non-primed (targeted) strand with a clear gradient centered at the priming site (Richter, Dy et al. 2014, Staals, Jackson et al. 2016). A “sliding” model has been proposed to explain these patterns: the spacer acquisition machinery (including the Cas1-Cas2 complex) is recruited to the targeted sequence and subsequently slides away from the priming site in a 3′ to 5′ direction preferentially on one strand and stops at an appropriate PAM site for spacer extraction (Heler, Marraffini et al. 2014). The translocation directionality is consistent with the helicase activity of Cas3 (Mulepati and Bailey 2013, Sinkunas, Gasiunas et al. 2013), indicating that Cas3 may travel in complex with Cas1-Cas2, and this notion is supported by single-molecule imaging (Redding, Sternberg et al. 2015, Wright, Nunez et al. 2016). Besides the sliding model that describes the overall patterns, another model regarding the molecular basis of spacer extraction suggests that double-stranded Cas3 degradation products are preferentially used as donors for Cas1-Cas2 (Swarts, Mosterd et al. 2012, Kunne, Kieper et al. 2016, Severinov, Ispolatov et al. 2016). These two models are not necessarily mutually exclusive, as close interactions between the interference and adaptation machineries are likely involved in robust CRISPR adaptation (Babu, Beloglazova et al. 2011, Richter, Gristwood et al. 2012, Sternberg, Richter et al. 2016). It is possible that, depending on if Cas3 is activated (by some as-yet-unclear signal) in its nuclease activity, the processing of targeted DNA could contribute to spacer acquisition through either Cas1-Cas2-Cas3 co-sliding (the sliding model) or Cas3 degradation followed by Cas1-Cas2 recycling for protospacer extraction (the alternative model).

Despite the accumulated knowledge on primed adaptation, a number of factors limit direct comparison between most of the previous studies. Specifically, due to the high interference efficiency against *bona fide* targets, most studies used mismatched priming sequences with either a non-canonical PAM or mutations in the seed sequence of a protospacer (Datsenko, Pougach et al. 2012, Savitskaya, Semenova et al. 2013, Fineran, Gerritzen et al. 2014, Li, Wang et al. 2014, Richter, Dy et al. 2014, Staals, Jackson et al. 2016). Such target mismatches not only affect interference (Wiedenheft, van Duijn et al. 2011, Xue, Seetharam et al. 2015), but influence the efficiency of primed adaptation (Fineran, Gerritzen et al. 2014, Li, Wang et al. 2014, Kunne, Kieper et al. 2016, Xue, Whitis et al. 2016), and may pose an impact on how spacers are acquired during priming (Redding, Sternberg et al. 2015, Vorontsova, Datsenko et al. 2015). To prime with a *bona fide* target, others used either inducible expression or anti-CRISPR regulated systems to control interference (Vorontsova, Datsenko et al. 2015, Semenova, Savitskaya et al. 2016, Staals, Jackson et al. 2016). In fact, these *bona fide* targets, despite being cleaved rapidly, were shown to be capable of inducing CRISPR adaptation with an even higher efficiency (Xue, Seetharam et al. 2015, Semenova, Savitskaya et al. 2016, Staals, Jackson et al. 2016). To avoid these complicating factors, the focus of the current study is on a relatively interference-permissive, type I-C CRISPR-Cas system in *L. pneumophila* -a system which is ideally suited to the study of priming with a *bona fide*, perfect-match target. Along with this inherent experimental strength, type I-C systems remain relatively understudied, despite representing the second most abundant type of CRISPR-Cas systems in prokaryotes (Makarova, Haft et al. 2011, Makarova, Wolf et al. 2015).

## Results

### Priming of the permissive *L. pneumophila* type I-C CRISPR-Cas induces robust spacer acquisition

In our previous work, we experimentally showed that a perfectly-targeted plasmid can temporarily co-exist, without detectable mutations (in either plasmid or CRISPR-Cas locus), with the type I-C CRISPR-Cas system in *L. pneumophila* str. Toronto-2005 (Rao, Guyard et al. 2016). These escaped transformants displayed a gradual plasmid loss during non-selective axenic passages and clear spacer acquisition events induced at the end (Rao, Guyard et al. 2016). Here we exploited this robust adaptation system to study spacer acquisition in the type I-C system in depth (Fig. 1A). A targeted plasmid that includes the CRISPR spacer 1 (Sp1) sequence and a canonical TTC PAM (Mojica, Diez-Villasenor et al. 2009, Leenay, Maksimchuk et al. 2016) on either the plus strand (pSp1(+)) or minus strand (pSp1(–)) was used to prime spacer acquisition (we refer to protospacer as the target identical to the spacer sequence and PAM as the 5′-3′ sequence upstream of the protospacer on the untargeted strand). These targeted plasmids showed a ∼1% relative transformation efficiency compared with untargeted control plasmids, and the escaped transformants were passaged without antibiotic selection for 15 generations to induce spacer acquisition events that we subsequently cataloged by PCR amplification, gel extraction and deep sequencing (Fig. 1A). Around 2 million new spacers were extracted from Illumina raw reads in each priming experiment, and mapped to potential sources including the priming plasmid or the bacterial chromosome. The vast majority (>99.7%) of spacers were derived from the plasmid, with the remaining few from the chromosome or unknown sources (possibly due to chimeric sequences or sequencing errors; Fig. 1B). Collectively these numerous spacer sequences covered all available TTC canonical PAM sites on the plasmid (Table S1), suggesting a sufficient sequencing depth to represent the CRISPR-adapted population.

**Figure 1:**
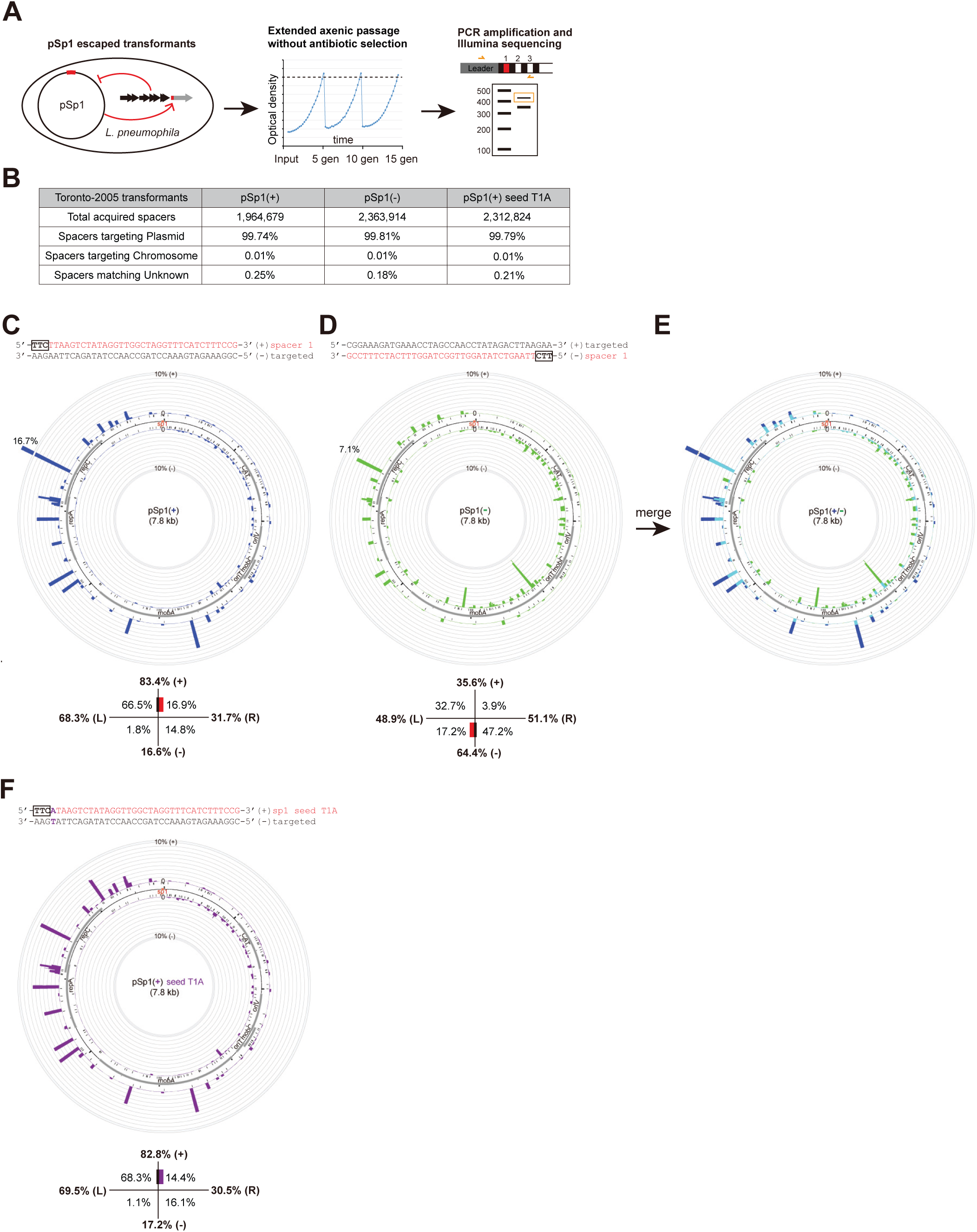
Primed spacer acquisition by *L. pneumophila* type I-C CRISPR-Cas occurs in a strand-biased manner. **A.** Schematic workflow to characterize primed spacer acquisition. Escaped transformants of targeted plasmids were passaged for 15 generations without antibiotic selection to enrich for spacer acquisition. CRISPR loci were PCR amplified and adapted arrays were further isolated through gel size selection. Amplicons were subjected to Illumina sequencing, and acquired spacers were extracted from raw reads and mapped to either the plasmid or the bacterial chromosome. **B.** The vast majority of acquired spacers during priming were derived from the plasmid instead of the chromosome. **C-D.** Circos plots of acquired spacers mapped to the pSp1 priming plasmid where the priming protospacer (identical to the spacer 1 sequence from the type I-C system) is either on the plus (+) strand (**C**) or on the minus (–) strand (**D**). In the strand-specific mappings, bars protruding inside and outside of plasmid circle represent spacers matching the minus and plus strand of the plasmid, respectively, and the height of bars indicates the number of spacers mapped to indicated positions. Note that a secondary scale was used for plasmid loci acquired at a frequency of over 10% of all spacers. The frequency of the major spacer acquisition hotspot is indicated. To numerically represent the overall spacer acquisition patterns, the plasmid is divided into four geographic fractions relative to the priming protospacer (denoted by the colored rectangle): the 5′ half (Left) and the 3′ half (Right) on the + strand, and the 3′ half (Left) and the 5′ half (Right) on the – strand. **E.** A merged view of the two mappings was created where overlapped coverages were shown in cyan. **F.** Priming by an imperfect target with a seed mismatch showed similar overall patterns of spacer acquisition as priming using a *bona fide* target. Each Circos plot in the figure represents the average of two independent biological replicates.

### Primed spacer acquisition occurs in a strand-biased fashion but is influenced by local hotspots

Through Sanger sequencing of 23 acquired spacers, we previously observed a moderate preference (74%) of spacers derived from the same strand as the priming protospacer (Rao, Guyard et al. 2016). With a much higher sequencing depth, we comprehensively re-examined the patterns of spacer acquisition from the plasmid. When the priming protospacer is on the plus strand of the plasmid, a majority (83%) of spacers are mapped to the same strand, and an obvious enrichment of acquisitions is seen from the 5′ region proximal to the priming site on both strands (Fig. 1C). When the priming sequence is flipped to the minus strand, the preferred strand of acquisition is also switched, with 64% spacers derived from the minus strand, and as before, more spacers are mapped to the 5′ region of the priming site relative to the 3′ region (Fig. 1D). A clear correlation between the directionality of the priming sequence and the strand preference of acquired spacers is shown from the merged view of pSp1(+) and pSp1(–) mappings (Fig. 1E). These observations are consistent with a strand-specific 3′ to 5′ translocation of Cas3 starting from the priming site.

Besides the strand bias and positional gradient, we observed a wide range of acquisition frequencies across the plasmid. Among all 238 TTC PAM sites, 30 positions each accounted for >1% of all acquisitions in at least one priming experiment, and 62 were acquired at <0.05% frequencies in both priming settings (Table S1). Strikingly, we identified one locus in the coding strand of *repC* that consistently ranked as one of the most frequently acquired spacers regardless of the primed strand (Fig. 1C, D). Interestingly, we did not observe an obvious enrichment of spacers from the origin of plasmid replication (oriV) and open reading frames (*cat*, *mobC*, etc.) - known hotspots in naïve adaptation and primed adaptations in other type I systems (Levy, Goren et al. 2015, Vorontsova, Datsenko et al. 2015, Staals, Jackson et al. 2016).

Lastly, we examined if a mismatched protospacer primes the type I-C system differently from a perfect match. While a single T1A mutation in the seed sequence increased the plasmid relative transformation efficiency from ∼1% to ∼9%, the overall patterns (strand bias and positional gradient) of acquired spacers were largely unchanged (Fig. 1C, F). These data are consistent with models in which primed spacer acquisition using perfect (interference-driven) or imperfect matches involves shared molecular mechanisms (Semenova, Savitskaya et al. 2016, Staals, Jackson et al. 2016).

### Sequence specificity contributes to the acquisition hotspot

To identify the factors contributing to the high acquisition efficiency of the hotspot, we first examined the hotspot region for the presence of any outstanding feature: PAM density, GC content, origin of replication or predicted small RNA transcription. In the absence of an obvious signal from any of these features, we hypothesized that some other sequence specificity of the hotspot region underlies the high acquisition efficiency. Thus, we generated a set of plasmids carrying mutations upstream, downstream, or within the hotspot while maintaining the *repC* codon to avoid any side-effects due to amino acid changes (Fig. 2A).

**Figure 2:**
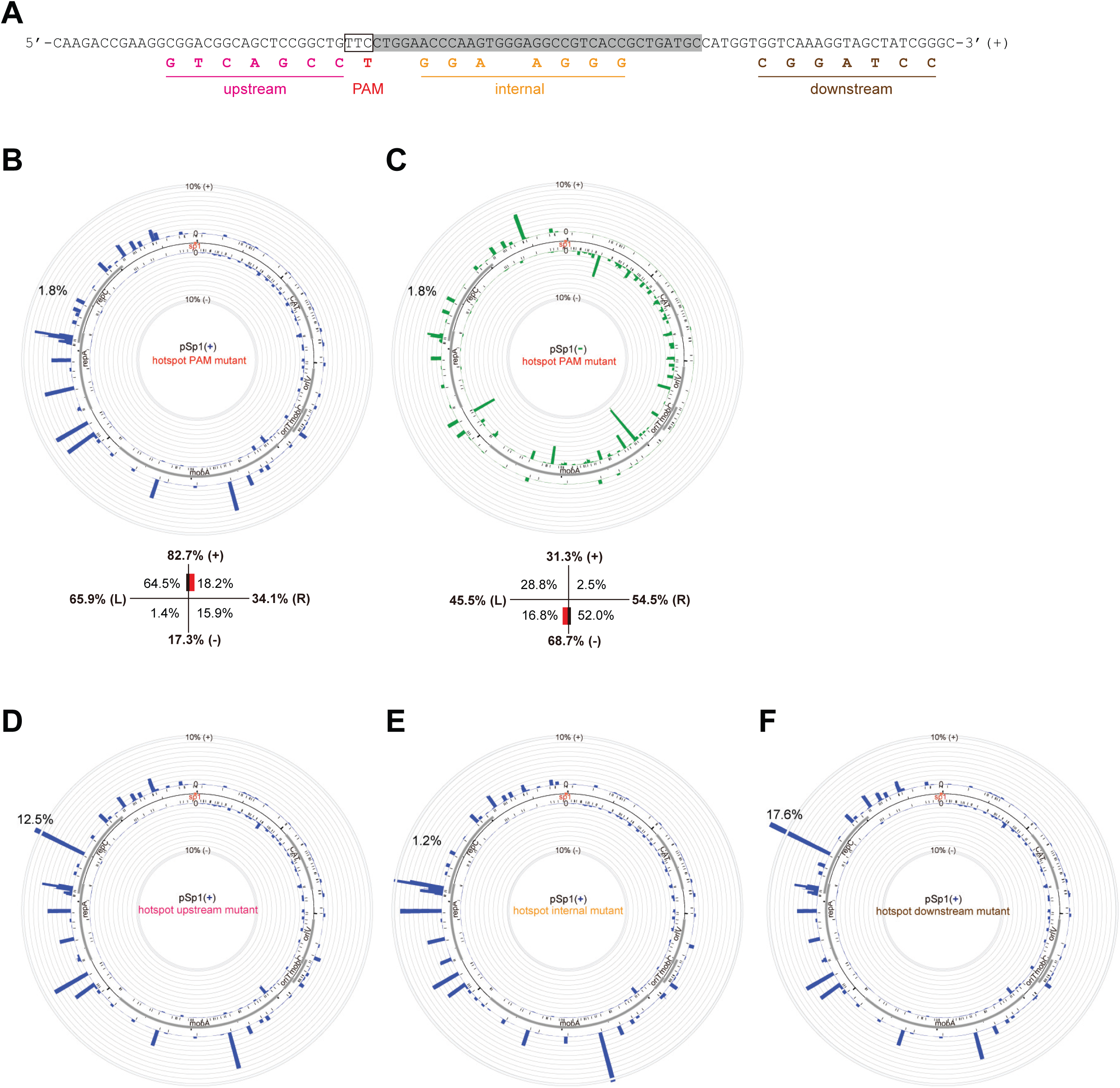
The spacer acquisition hotspot is reliant on its internal sequence. **A.** Mutations were introduced, with *repC* codons maintained, at the upstream, PAM, internal or downstream sequences of the major spacer acquisition hotspot to examine factors contributing to the high acquisition frequency. **B-C.** The mutation at the PAM dramatically reduced the acquisition frequency at the hotspot, while the overall patterns of spacer acquisition remained largely unaffected. Note that the mutation does not eliminate available PAM, but shifted the PAM 1 nt away. **D-F.** Mutations within, but not flanking, the hotspot also largely decreased the acquisition frequency at the hotspot. Each Circos plot in the figure represents the average of two independent biological replicates.

We first tested the PAM mutant in pSp1(+) where the TTC PAM of the hotspot is changed to TTT (where coincidentally another TTC motif is made with +1 nt shift). The acquisition efficiency of the mutant sequence decreased by 9-fold (16.7% to 1.8%) (Fig. 2B). We also observed a large reduction of acquisition efficiency at the hotspot by introducing the same mutation in pSp1(–) (Fig. 2C). Importantly, by comparing these hotspot PAM mutants with the *wild type* plasmids, we did not observe a major difference in spacer acquisition patterns due to the elimination of the hotspot, i.e. the imperfect mirroring of strand bias (>80% plus in pSp1(+) and <70% minus in pSp1(–)) is retained. This observation suggests that the local hotspot likely does not affect the initial sliding of the adaptation machinery whose strand preference is partially skewed to the plus strand due to unknown factors.

We next examined the other regions of the hotspot by introducing different sets of mutations in pSp1(+). The acquisition efficiency of the hotspot was dramatically eliminated by internal mutations, slightly reduced by changes upstream, and not reduced at all by the downstream perturbations (Fig. 2D-F). Elimination of the hotspot increases acquisition frequencies of other plasmid loci (Fig. 2B, E), suggesting that its loss modifies the availability of the adaptation machinery to other loci. The major impacts of the PAM mutation and the internal substitutions suggest that some sequence specificity within the hotspot, likely in the 5′ end, contributes to the acquisition preference at this locus.

### Analysis of acquired spacers reveals an alternative PAM and extensive acquisition inaccuracies

Of all acquired spacers from the plasmid, those having a TTC PAM account for 92.5% and 90.0% in the pSp1(+) and pSp1(–) priming experiment, respectively (Fig. 3B). We examined the trinucleotide sequence upstream of all acquired spacers from pSp1(+) for the abundance of other PAMs (Fig. S1A). The ∼2 million acquired spacers from pSp1(+) are next to 2,978 different PAM loci. While most (90%) of these PAM loci were acquired rarely (with <0.01% frequencies), some PAM sequences other than TTC were oversampled, suggesting one or more alternative PAMs. Based on the frequency rankings of the trinucleotide PAM sequences, the second most frequent PAM is TTT, followed by four TCN motifs (Fig. S1A). As these less frequent PAMs share a 2 nt identity with TTC, which could derive from slipping events where the real PAM is still TTC located nearby (Shmakov, Savitskaya et al. 2014), we first suspected that the TTT and TCN PAMs might be due to –1 nt slips (upstream) and +1 nt slips (downstream), respectively (Fig. 3A). Thus, we separately reanalyzed acquired loci with each of these PAMs with respect to their flanking sequence. Spacers with a TCN PAM showed a major T signal further upstream, consistent with +1 nt slips from TTC (Fig. 3C). However, spacers with a TTT PAM did not show an outstanding signal next to the trinucleotide, suggesting that TTT is an alternative PAM other than TTC (Fig. 3C). Consistent with this interpretation, acquired spacers with a TTT PAM showed independent localizations relative to those with a TTC PAM (Fig. S1B).

**Figure 3:**
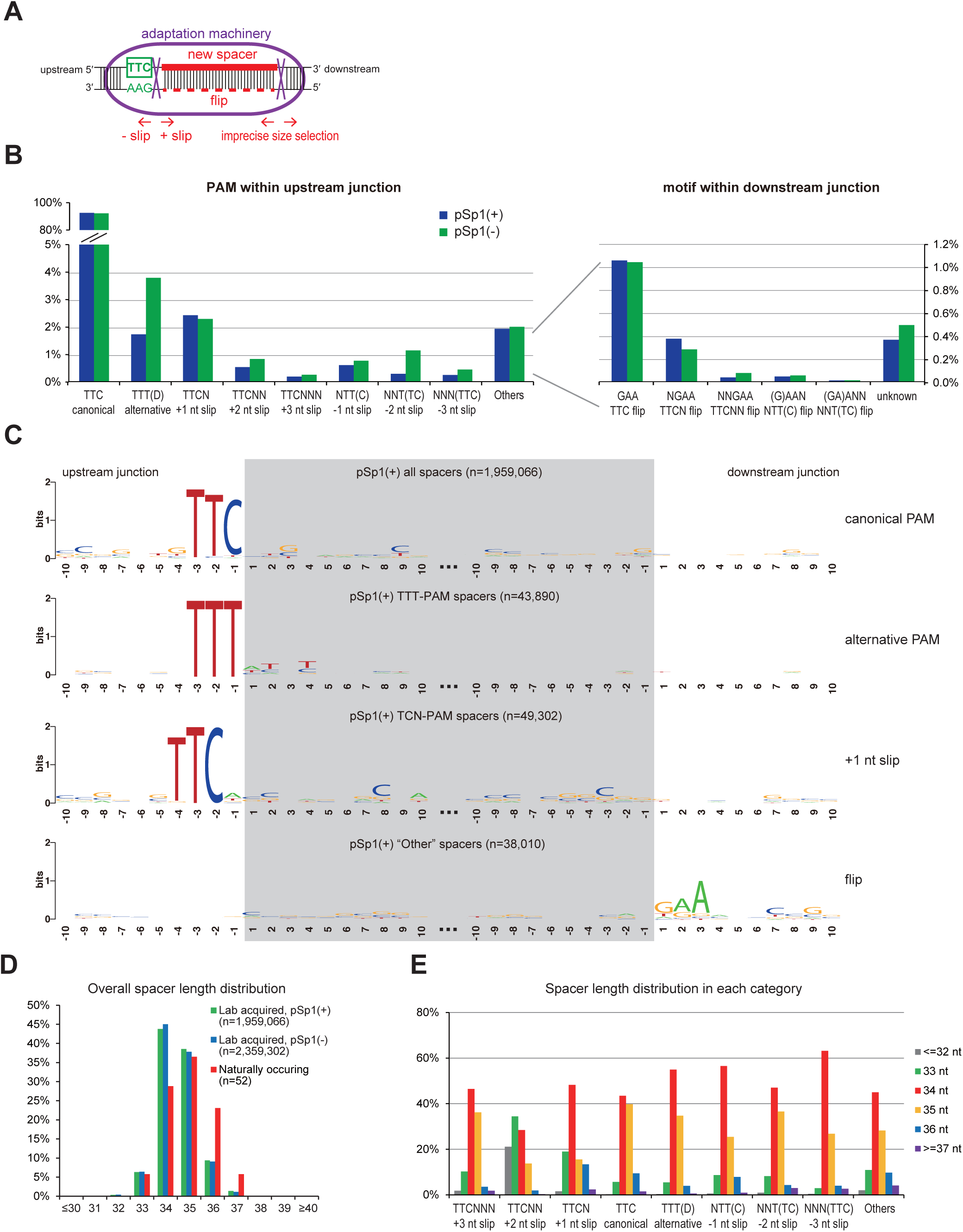
PAM preference and acquisition inaccuracies in primed adaptation. **A.** Schematic representation of spacer selection by the adaptation machinery. In most cases, the machinery extracts the double-stranded sequence immediately downstream of the PAM, with an inexact molecular ruler at the PAM-distal end. Less frequently, the machinery shifts a few nucleotides downstream (+ slip) or upstream (– slip) at the PAM-proximal end, causing slipping events. Flipping events were also observed where the double-stranded DNA substrates were incorporated in an opposite orientation into the CRISPR array. **B.** Based on the PAM localization within the upstream or downstream junction, spacer acquisition events were categorized into different types, with their frequencies shown for both pSp1(+) and pSp1(–) priming. Note that for the alternative TTT PAM, spacers with the first nucleotide being C were excluded as these were classified as potential -1 nt slipping events. **C.** Sequence Logo of the upstream and downstream 20 nt junctions of indicated categories of spacer acquisitions from pSp1(+). **D.** Length distribution of all acquired spacers primed by pSp1(+) and pSp1(–), compared with the native spacers from type I-C CRISPR loci in *L. pneumophila* ST222 strains. **E.** Length distribution of acquired spacers from each slipping category.

As we observed +1 nt slips, we wondered if there were other types of acquisition errors. We systematically examined + slips where the acquisition machinery extracts protospacers further downstream at the PAM and – slips where cleavage happens further upstream (Fig. 3A). Indeed, besides +1 nt slips that occurs at a ∼2% frequency, other types of slips do happen - though at 0.1%∼1% frequencies, with a decreasing trend as the slipping goes further (Fig. 3B). Apart from the aforementioned classes of spacer acquisition where the upstream sequence of the plasmid contains either a TTC or a TTT PAM, ∼2% spacers remained unexplained. When we examined the target sequence upstream and downstream of these spacers, we observed a clear GAA signal directly downstream (Fig. 3C). This downstream GAA signal is consistent with a phenomenon known as “flipping” - where a double-stranded DNA pre-spacer is extracted next to a TTC PAM but subsequently integrated in an opposite direction into the CRISPR array so that the reverse complementary strand is used as spacer (Shmakov, Savitskaya et al. 2014). Indeed, we identified ∼1% spacers derived from potential flips with an original TTC PAM, 0.3%∼0.4% spacers from a combination of flips and +1 nt slips, and even rarer still - combinations of flips and other types of slips (Fig. 3B). When combined, the canonical TTC PAM, the alternative TTT PAM and slipping and flipping events explain >99.5% of all acquired spacers from the priming plasmid (Fig. 3B).

The native spacers in *L. pneumophila* type I-C CRISPR arrays range from 33 to 37 nt, with 35 nt being the most frequent length. We next asked if acquired spacers had a similar length distribution. Compared with native spacers, laboratory acquired spacers showed a slight shift towards shorter lengths, with 34 nt being the most frequent (Fig. 3D). Compared with type I-E and type I-F systems that acquire spacers mostly (∼90%) with a uniform length (Datsenko, Pougach et al. 2012, Savitskaya, Semenova et al. 2013, Fineran, Gerritzen et al. 2014, Richter, Dy et al. 2014, Staals, Jackson et al. 2016), the type I-C system acquired spacers with a broad range of lengths - suggesting a remarkably imprecise molecular ruler in the adaptation machinery. This inaccuracy is largely unattributed to either slipping or flipping events, as spacers with a canonical TTC PAM showed a similarly large distribution of length (Fig. 3E). Interestingly, in +1 nt slips and +2 nt slips, we observed a distinct distribution of spacer length, with an increasing preference for shorter (≤33 nt) ones (Fig. 3E). This is in contrast with the observations in the *P. atrosepticum* type I-F system where – slips instead of + slips correlated with aberrant spacer lengths (Staals, Jackson et al. 2016), indicating another molecular distinction between these two adaptation machineries. Taken together, we identified extensive acquisition inaccuracies in the *L. pneumophila* type I-C system. A representative example of these inaccuracies can also be found at the major spacer acquisition hotspot (Fig. S1C).

### Systematic quantification of interference efficiencies confirms a hierarchy of preferred PAMs

PAM recognition in spacer acquisition is attributed to the adaptation machinery - and this process is likely independent from the Cascade interference complex that by itself recognizes the PAM and binds to target. To examine the possible co-evolution of PAM recognition by these two machineries (Kunne, Kieper et al. 2016), we asked if the Cascade interference complex also recognizes alternative PAMs other than the canonical TTC motif. Indeed, using an *in vivo* positive screen, Leenay *et al*. recently identified TTC, CTC, TCC and TTT, with decreasing preferences, as functional PAMs for interference in the *Bacillus halodurans* type I-C system (Leenay, Maksimchuk et al. 2016). Here, we performed a plasmid-removal based screen to examine functional PAMs for interference in *L. pneumophila* (Fig. 4A). We transformed a plasmid library containing a full spacer 1 match with a randomized trinucleotide PAM into either *L. pneumophila* str. Toronto-2005 *wild type* or Δ*cas3*. By analyzing the PAM abundance in the survived plasmid pools using high-throughput sequencing, we identified PAMs that were depleted to different degrees by the *wild type* type I-C system. Among the 64 PAM sequences, TTC achieved the highest protection efficiency of >99.9%, 6 others (TTT, CTT, CTC, TTA, TTG and TCC) within the range of 95% ∼ 99.5%, and 11 more above 50% (Fig. 4B). It is noteworthy that TTT is the second most interference-efficient PAM, consistent with our observation that TTT is also the second most frequent PAM used in spacer acquisition. Many of the less protective PAMs share a 2 nt identity with TTC, suggesting that a 1 nt perturbation of the PAM would still allow some functionality. We confirmed the observed hierarchy of PAM activities using a CFU-based plasmid transformation efficiency assays of 8 selected PAMs (Fig. 4C). Inspection of the escaped transformants of TTT PAM plasmids showed spacer acquisition events similar to the transformants of the *bona fide* target and mismatched protospacer plasmids, suggesting that the alternative PAM primes in a similar manner (Fig. 4D). Together, our assay suggests that the *L. pneumophila* type I-C system possesses a broader range of active PAMs in interference than in adaptation.

**Figure 4:**
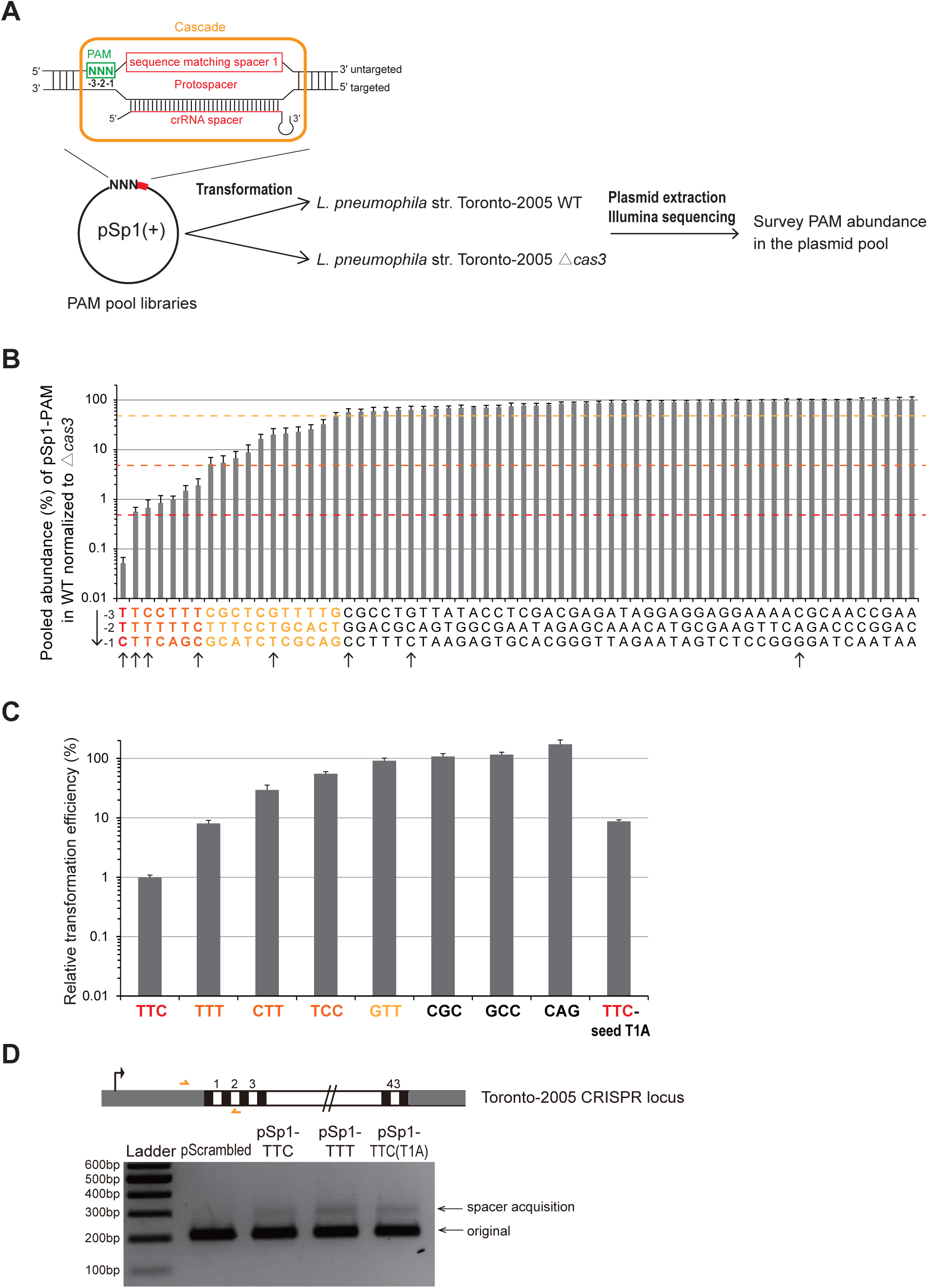
PAM preference for the *L. pneumophila* type I-C CRISPR-Cas interference. **A.** Schematic workflow to characterize functional PAMs for CRISPR interference. A pool of plasmids containing the spacer 1 sequence and a random trinucleotide PAM was generated and transformed into either *L. pneumophila* str. Toronto-2005 *wild type* or Δ*cas3*, and the abundance of each PAM sequence in the pool was quantified through Illumina sequencing. **B.** Pooled abundances were derived by normalizing the ratio of each PAM in the *wild type* transformant pool to that in the Δ*cas3* pool. These relative abundances categorized PAM sequences into different preferences for CRISPR interference. **C.** Individual plasmid transformation efficiency assay confirmed, with a lower sensitivity, the observations in the Illumina-based pooled assay. Plasmids containing either spacer 1 and an indicated PAM or a scrambled control sequence were electroporated into *L. pneumophila* str. Toronto-2005 *wild type*. The relative transformation efficiency is calculated by normalizing transformation efficiency of the spacer 1 plasmids to that of the control plasmid. Error bars represent the SEM of three biological replicates. **D.** Spacer acquisition was also observed in escaped transformants of targeted plasmids with the alternative TTT PAM.

### Truncation of the type I-C array leads to a dramatic increase in interference and frequent spacer loss

As *L. pneumophila* type I-C CRISPR-Cas is relatively permissive for interference, we wondered how spacer acquisition efficiency would change if target cleaving by this system is made more efficient. While studying a minimized type I-C array that contains only a single spacer, we made an unexpected observation that allowed us to further explore the relationship between interference efficiency and spacer acquisition. We generated a CRISPR array-minimized strain in which all 43 spacers except the spacer 1 were deleted in *L. pneumophila* str. Toronto-2005. Remarkably, the sole spacer in this strain showed a ∼100-fold increased protection efficiency against its matching protospacer as compared to the parental strain that contains a full-length (43 spacers) array (Fig. 5A). When examining the CRISPR loci in the less frequent escaped transformants, no spacer acquisition was observed, in contrast to what we observed for the more permissive, full-length array strain. Instead, these escaped transformants showed clear spacer loss events (Fig. 5A). To test if the modified CRISPR array is adaptable, we next transformed the spacer 1 only strain with a mismatched target plasmid (carrying a T1A seed mutation). Use of this mismatched target led to both decreased interference efficiency and robust spacer acquisition, demonstrating the adaptability of the minimized CRISPR array under certain conditions (Fig. 5A). Consistent with the observations using agarose gel, by quantifying spacer dynamics in the single-spacer strain transformants of the spacer 1 targeted plasmid, we observed 39% spacer loss frequency in the CRISPR loci and a ∼100-fold lower spacer acquisition frequency relative to the *wild type* strain transformants (Fig. 5B). As the minimized array was designed to contain two similar but distinct repeat sequences flanking spacer 1, we examined the spacer loss events and found that most loci retained the downstream repeat, consistent with a mechanism of homologous recombination (Fig. 5C). It is also noteworthy that we quantified spacer dynamics in the single-spacer strain transformed with an untargeted plasmid. We observed a detectable <0.1% spacer loss frequency without spacer targeting, suggesting that spacer loss events naturally occur in CRISPR loci at a low frequency and can be enriched under selection (Fig. 5B).

**Figure 5:**
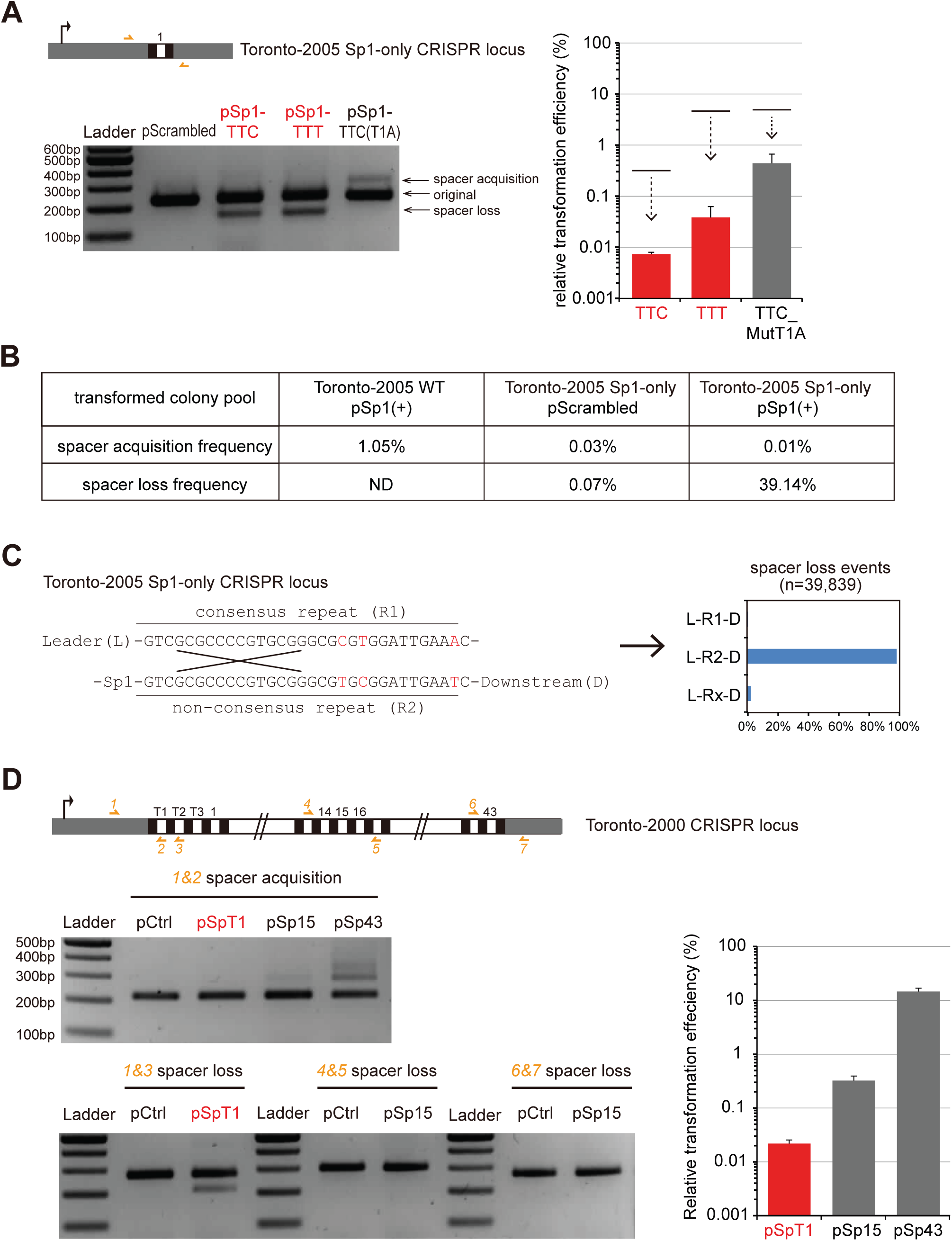
Highly-efficient interference leads to spacer loss rather than spacer acquisition. **A.** Spacer loss, rather than spacer acquisition, was seen in escaped transformants when the array-minimized (Sp1-only) CRISPR-Cas system highly efficiently (>99.9% by relative transformation efficiency) protects against targeted plasmids. Plasmid transformation efficiency assay was performed to measure interference efficiencies of the modified CRISPR-Cas system (compared with those of the original system, shown by the upper lines). The resulting transformants were examined by PCR amplification for the dynamics of CRISPR loci. **B.** Quantification of spacer acquisition and spacer loss frequencies in plasmid transformants (colonies pool without further passages) of *L. pneumophila* str. Toronto-2005 *wild type* or Sp1-only. Spacer loss frequency is not determined for *wild type* transformants because only the leader-end of the CRISPR array was surveyed by PCR. **C.** Most spacer loss events retained the downstream non-consensus repeat (R2), shown in the bar graph, consistent with a mechanism of homologous recombination between the two flanking repeats (R1 and R2). The few Rx repeats contain mismatches to both R1 and R2 and may derive from sequencing errors. **D.** Spacer loss was also observed in another native type I-C CRISPR-Cas system in *L. pneumophila* str. Toronto-2000 where the first spacer (SpT1) is highly efficient in interference. Plasmids containing a targeted sequence for one of the three indicated spacers showed different relative transformation efficiencies in *L. pneumophila* str. Toronto-2000. The resulting transformants were tested for spacer acquisition or spacer loss by PCR using indicated primers. Error bars represent the SEM of three biological replicates.

Our observations using the minimized array suggest that when CRISPR interference reaches a threshold of efficiency to no longer tolerate the temporary co-existence between the target and a functional CRISPR-Cas, the resulting transformants would select for spacer loss rather than spacer acquisition. To further explore the interference efficiency threshold between spacer acquisition and spacer loss, we took advantage of a naturally occuring type I-C system with a greater protection. The type I-C CRISPR-Cas system in *L. pneumophila* str. Toronto-2000 differs from the one in *L. pneumophila* str. Toronto-2005 by three newly acquired spacers (Rao, Guyard et al. 2016). Transformation of a plasmid targeted by the first spacer T1 in this array, resulted in a very low transformation efficiency and correspondingly spacer loss instead of spacer acquisition in the rare escaped transformants (Fig. 5D). Examination of downstream spacers in this system, which provide less efficient protection, suggests that less efficient interference is correlated with stronger spacer acquisition (Fig. 5D). These data, together with the observations in the single-spacer strain, indicate that an interference efficiency of >99.9% by plasmid transformation is a good empirical indicator for an absence of primed adaptation using *bona fide* targets in the *L. pneumophila* type I-C system.

## Discussion

A clear interplay between CRISPR interference and adaptation has been established (Sternberg, Richter et al. 2016, Wright, Nunez et al. 2016). For CRISPR-Cas systems that execute the interference very efficiently (interference-strict), a slow or delayed target degradation (by target mismatches or other means) is often necessary to achieve an efficient primed adaptation (Kunne, Kieper et al. 2016, Semenova, Savitskaya et al. 2016). Here we confirmed the observation using a native interference-permissive system. We propose that when the cleaving efficiency of a system allows the temporary coexistence of target and a functional CRISPR-Cas, robust spacer acquisition predominates. In contrast, when this system is made highly efficient in spacer targeting, spacer loss events were selected for and the resulting transformants showed a lack of primed adaptation. The observed low frequency of natural spacer loss in the CRISPR array also provides another perspective of spacer dynamics - that CRISPR-Cas systems can readily update and diversify its immunological memory through both spacer acquisition and spacer loss, thus providing raw materials of evolution.

Our analyses of spacer acquisition patterns (Fig. 6) are consistent with the sliding model in which the adaptation machinery (which includes Cas1-Cas2 and possibly Cas3) translocates 3′ to 5′ preferentially on the untargeted strand before stopping at an appropriate sequence to extract a protospacer for subsequent integration into the CRISPR array. Several similarities and differences exist between *L. pneumophila* type I-C acquisition and what has been described for other systems in other bacteria (Fig. 6). We observe similar spacer acquisition patterns between the *L. pneumophila* type I-C system and the *H. hispanica* type I-B system (Li, Wang et al. 2014): both systems show a moderate (70∼80%) overall bias towards the untargeted strand and a positional preference for the 5′ region on both strands relative to the priming site. In contrast, the *P. atrosepticum* type I-F system prefers spacers on the targeted strand and samples spacers in a narrower distance from the priming sequence (Richter, Dy et al. 2014, Staals, Jackson et al. 2016). Type I-F's opposite strand preference could possibly be due to an opposite spatial organization of the adaptation complex relative to the PAM recognition. The *E. coli* type I-E system, shows robust spacer acquisitions from the untargeted strand, like type I-C and type I-B, but proximity to the priming sequence appears to have little influence on the overall pattern of spacer acquisition (Datsenko, Pougach et al. 2012, Savitskaya, Semenova et al. 2013, Fineran, Gerritzen et al. 2014). To explain the discrepancies of positional preference (sampling distance) in different systems, a variable processivity of Cas3 in different systems was proposed (Redding, Sternberg et al. 2015). This model would suggest that the type I-C and type I-B systems should have an intermediate level of Cas3 processivity (between a highly processive type I-E system and a less processive type I-F system). The different levels of strand bias in these systems, on the other hand, may attribute to different degrees of “PAM-independent processing” in which Cas3 is recruited with the help of Cas1-Cas2 and travels bidirectionally (Redding, Sternberg et al. 2015).

**Figure 6:**
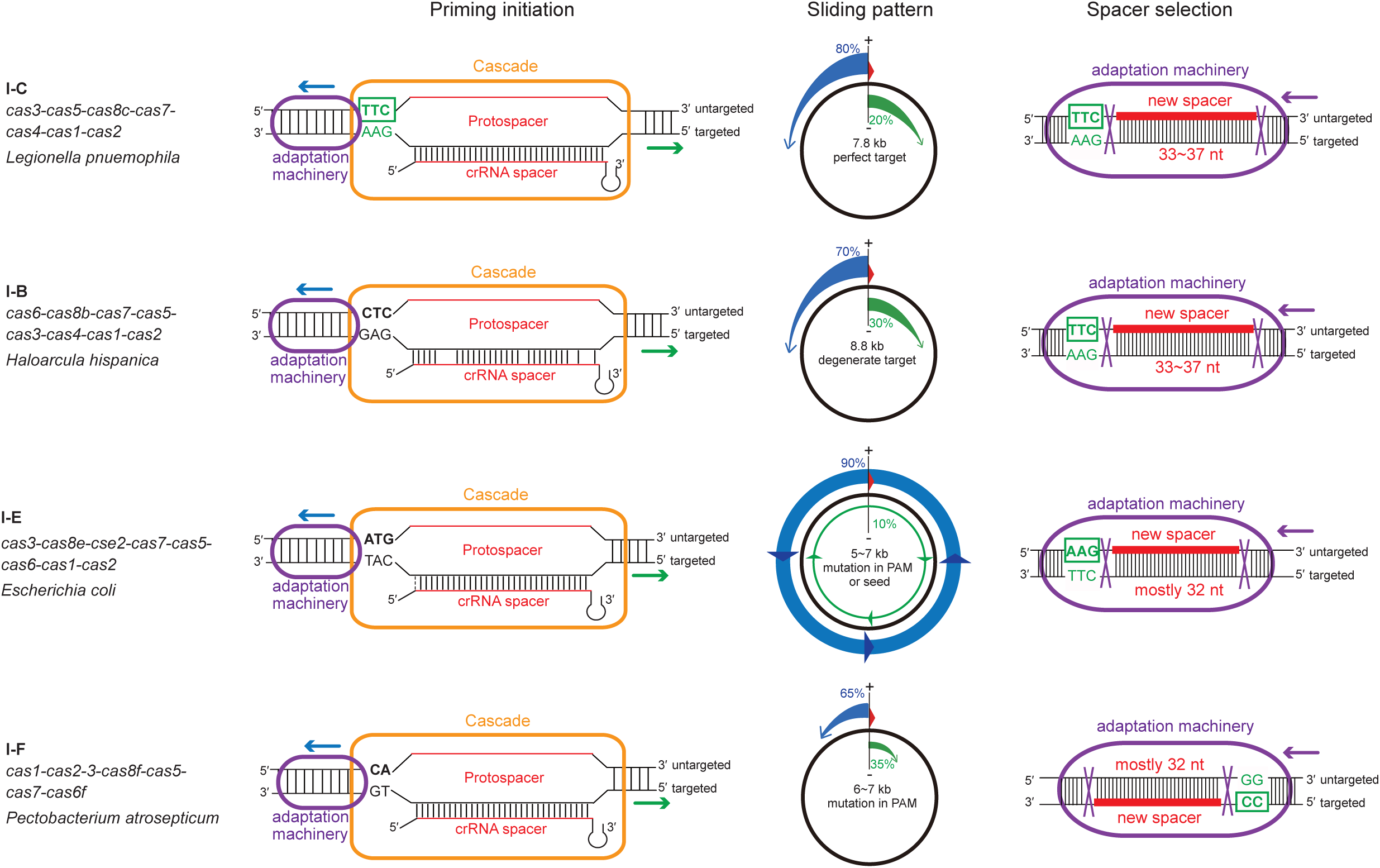
Schematic summary of primed spacer acquisition in type I CRISPR-Cas. Primed spacer sampling is separated into three steps: 1) priming initiation where the Cascade-crRNA complex binds to the targeted DNA and recruits the adaptation machinery; 2) the adaptation complex surveying the plasmid, consistent with a 3′ to 5′ sliding with variable strand specificity; 3) spacer selection from another plasmid locus by the adaptation complex upon recognition of an appropriate PAM sequence. Comparisons of each step in the type I-C system versus type I-B (Li, Wang et al. 2014), type I-E (Datsenko, Pougach et al. 2012, Savitskaya, Semenova et al. 2013, Fineran, Gerritzen et al. 2014) and type I-F systems (Richter, Dy et al. 2014, Staals, Jackson et al. 2016), show similarities and distinctions in molecular mechanisms. Note that most previous studies used an imperfectly-targeted priming sequence with mutations in either the PAM or the protospacer sequence.

Compared with the earlier study of the *H. hispanica* type I-B system that used Sanger sequencing (Li, Wang et al. 2014), we have achieved a much higher survey depth through next-generation sequencing, thus enabling a more comprehensive examination of spacer acquisition details. In PAM preference, in addition to the canonical TTC PAM (∼90% of all acquired spacers), we identified an alternative TTT PAM (2∼4%) and extensive slipping and flipping events (6∼8%). With respect to spacer size selection, we observed a flexible choice more similar to the type I-B system (Li, Wang et al. 2014) than the highly stringent type I-E (Savitskaya, Semenova et al. 2013, Fineran, Gerritzen et al. 2014) and type I-F (Richter, Dy et al. 2014, Staals, Jackson et al. 2016) systems. These observations suggest that the type I-C Cas1-Cas2 protein complex is relatively promiscuous in spacer extraction from pre-spacer substrates. The similarity between the type I-C and type I-B systems is also consistent with their closer Cas1-based phylogenetic relationship relative to the other two systems (Fig. S3). Further insights into the molecular basis of spacer acquisition stringency may be derived from a detailed structure based comparison of Cas1 and Cas2 from each system.

It is known that different CRISPR-Cas systems, as well as different spacers within one array, often show a wide range of interference efficiencies (Marraffini and Sontheimer 2008, Bikard, Hatoum-Aslan et al. 2012, Cady, Bondy-Denomy et al. 2012, Li, Wang et al. 2014, Xue, Seetharam et al. 2015, Qiu, Wang et al. 2016, Rao, Guyard et al. 2016) (Fig. 5D). Both technical and biological factors could contribute to this variation. On the one hand, transformation methods, plasmid copy number, bacterial culture conditions, etc. could all affect how efficiently invasive DNA is cleaved (Majsec, Bolt et al. 2016, Rao, Guyard et al. 2016, Severinov, Ispolatov et al. 2016). On the other hand, innate factors could also influence interference efficiency – such as expression levels of Cas proteins, transcription and processing efficiencies of individual spacers, and binding affinities between Cascade and crRNA (Xue, Seetharam et al. 2015, Hoyland-Kroghsbo, Paczkowski et al. 2016, Patterson, Jackson et al. 2016, Rao, Guyard et al. 2016). We observed a dramatic increase in interference efficiency for the same spacer (Sp1) when the CRISPR array was minimized. This could be due to a higher abundance of Sp1 crRNA, a relatively increased availability of Cas proteins for Sp1 (due to lack of competition with other spacers for loading), or a combination of both. Future experiments to examine the crRNA abundance and to over-express each Cas functional group (to determine limiting factors) will be necessary to test these hypotheses.

Consistent with previous studies of CRISPR adaptation (Paez-Espino, Morovic et al. 2013, Yosef, Shitrit et al. 2013), we observed a great range of spacer acquisition frequencies at different locations of the same element. While this variation could be affected by PAM specificity, strand specificity and strand-specific distance from the priming site, our examination of the major spacer acquisition hotspot points towards other factors that directly contribute to pre-spacer capture by the adaptation machinery. By introducing different mutations in the hotspot neighbourhood, we found that the internal sequence, but not the flanking nucleotides, contributes to the frequent acquisition. Based on these data, we speculate that some DNA motif or ssDNA secondary structure within the hotspot sequence (likely at the PAM-proximal end) could attract the adaptation machinery, and further systematic mutation experiments are required to identify the exact contributor. The intrinsic sequence specificity of the type I-C hotspot stands in stark contrast to a study of the *E. coli* type I-E system that showed a frequently-acquired protospacer was reliant on its upstream and downstream sequences (Yosef, Shitrit et al. 2013). Notably, we also did not observe a detectable enrichment of spacer acquisition from either the origin of plasmid replication or transcriptionally active regions - in contrast to what has been seen in type I-E and type I-F systems (Levy, Goren et al. 2015, Vorontsova, Datsenko et al. 2015, Staals, Jackson et al. 2016). These discrepancies further indicate a mechanistic distinction for pre-spacer capture in different systems and point towards the potential for diverse model systems to inform our understanding of the mechanisms underpinning CRISPR-Cas interference and adaptation.

Going forward, several features make the *L. pneumophila* type I-C system a good model system to study CRISPR-Cas functionality. First, type I-C systems represent one of the most common types of CRISPR-Cas systems yet nevertheless remain relatively understudied (Makarova, Haft et al. 2011, Makarova, Wolf et al. 2015). Second, our earlier comparative genomics data suggest that the system is naturally adaptable (Rao, Guyard et al. 2016). Third, the relatively permissive interference of the system allows the laboratory study of primed spacer acquisition within the context of perfectly matched target sequences. Lastly, based on our initial characterizations, the system displays several features that distinguish it from type I-E and type I-F systems, the two systems most exhaustively studied to date.

## Materials and Methods

### Bacterial strains and plasmids

*Legionella pneumophila* strain Toronto-2005 is a clinical isolate of Sequence Type 222 from Toronto, Canada, with a circularized genome available (Genbank CP012019) (Rao, Guyard et al. 2016). An RpsL^K43R^ streptomycin resistant derivate of the clinical isolate is used as *wild type* in this study. From this RpsL^K43R^ strain, a Δ*cas3* deletion mutant and an array-minimized (Sp1-only) strain were generated by allelic exchange as described (Ensminger, Yassin et al. 2012, Rao, Guyard et al. 2016). Specifically, in the Sp1-only strain, only the first repeat, Sp1 and the last repeat of the original array were retained. A closely-related ST222 strain, Toronto-2000, was also genome sequenced (Rao, Guyard et al. 2016). The priming plasmids were generated by cloning the insert (see Table S2) into the ApaI/PstI-cut pMMB207 backbone (Rao, Benhabib et al. 2013, Rao, Guyard et al. 2016) (see Supplemental File for the full pSp1(+) sequence). Our previous study using Illumina sequencing showed that this plasmid has an average copy number of 7.6 in *L. pneumophila* str. Philadelphila-1 (Rao, Benhabib et al. 2013). Site-directed mutagenesis (QuickChange II) was used to mutate the spacer acquisition hotspot in the original plasmid. Bacterial electroporation and axenic passage were performed as previously described (Rao, Guyard et al. 2016). After axenic passages for 15 generations, the CRISPR adaptation ratio in the bacterial population increased from ∼1% to ∼24%, as quantified by Illumina sequencing (data not shown). Each priming experiment was performed in two biological replicates and these replicates were largely consistent in spacer mappings (data not shown). Unless specified, data shown are averages of two replicates.

### PCR of CRISPR loci and preparation of Illumina libraries

Roughly 1 OD unit (∼1×10^9^) bacterial cells from either colony pool (containing at least 50 independent colonies) or axenic passage were used for genomic DNA extraction using the NucleoSpin Tissue kit (Machery-Nagel). CRISPR loci were amplified using the Kapa HiFi polymerase (Kapa Biosystems) and primers listed in Table S1. Raw PCR products of 20 amplification cycles were used for library preparation. In addition, to enrich for adapted CRISPR arrays, 30-cycle PCR products were concentrated by ethanol precipitation and separated in 6% acrylamide gel by running at 60V for 3 hours. A ∼70 bp higher band than the original array (∼350 bp) was extracted and DNA purified from the extraction was subjected to another 10-cycle PCR to increase the yield. These further size-selection steps to enrich for adapted arrays did not introduce significant bias relative to the raw PCR products (data not shown). Purified PCR amplicons were normalized by PicoGreen to 1 ng and processed using the Nextera XT kit (Illumina). Multiplexed libraries were subjected to Illumina NextSeq sequencing at 2 x 150bp read length (CAGEF, University of Toronto).

### Illumina reads processing and data analyses

Paired-end raw reads were first attempted to merge by FLASH (Magoc and Salzberg 2011) using “-m 50 -M 100 -x 0.02” settings. The unassembled single-end reads were quality trimmed by Trimmomatic (Bolger, Lohse et al. 2014) using “SLIDINGWINDOW:3:20 MINLEN:50” settings. These pre-processed reads were combined and processed using a Perl script (available upon request) to annotate the presence of leader sequence (L), CRISPR repeats (R), existing spacers (S), new spacers (X) and downstream sequence (D) in each read. The new spacers were extracted and aligned using blastn to either the priming plasmid or *L. pneumophila* str. Toronto-2005 genome. Blastn results were then summarized into coverages of each nucleotide in the plasmid and subjected to Circos visualization (Krzywinski, Schein et al. 2009). To examine the PAM preference, slipping and flipping of acquired spacers, flanking sequences of acquired spacers were extracted from the plasmid and subjected to Sequence Logo (Crooks, Hon et al. 2004) visualization. To avoid potential redundancy, flipping cases were only examined from spacers without a TTC or TTT PAM in the upstream junction. To quantify spacer acquisition and spacer loss frequencies, the following formulas were used, in which each item denotes the count of reads with the indicated annotation:

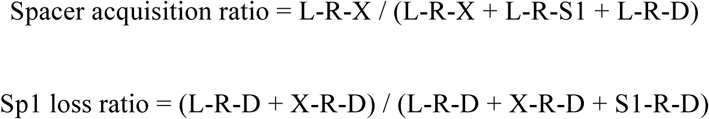

### Preparation and analyses of PAM plasmids pool

Oligos (see Table S2) with a randomized trinucleotide upstream of Sp1 sequence were annealed, digested and ligated into the ApaI/PstI-cut pMMB207 vector. A total of ∼3000 *E. coli* colonies were obtained after transformation and combined into a pool. Plasmids were extracted from the *E. coli* pool using the PureYield Plasmid Midiprep kit (Promega), and a control plasmid with a scrambled insert was spiked into the plasmid pool at ∼1% ratio. Roughly 1 µg of the pooled plasmids was electroporated into 4 OD units of *L. pneumophila* str. Toronto-2005 *wild type* or Δ*cas3* overnight culture. Three biological replicates of electroporation were performed. With 5 µg/ml chloramphenicol selection, over 3000 colonies were obtained from each electroporation. Plasmids were then extracted from these *L. pneumophila* transformants using the EZ-10 Spin Column Miniprep kit (Biobasic). Without any PCR amplification, these plasmid pools were subjected to the Nextera XT library preparation and Illumina NextSeq sequencing. After quality filtering, reads containing the Sp1 sequence (or the scrambled sequence) were extracted and PAM sequences were identified from these reads. PAM frequencies in *L. pneumophila* transformants were normalized to both the scrambled control and the *E. coli* plasmid pool.

### Data accessibility

The NextSeq sequencing data have been deposited in the NCBI Sequence Read Archive under the BioProject PRJNA360289.

## Acknowledgments

This work was supported by a Project Grant from the Canadian Institutes of Health Research (PHT-148819), the Connaught Fund (NR-2015-16), and an infrastructure grant from the Canada Foundation for Innovation and the Ontario Research Fund (30364) to AWE. The authors thank members of the Centre for the Analysis of Genome Evolution and Function (CAGEF) at the University of Toronto for performing Illumina sequencing. We thank members of the Ensminger laboratory for helpful discussions and for careful reading of the manuscript.

## Supplemental Information

**Supplemental Figure 1:**
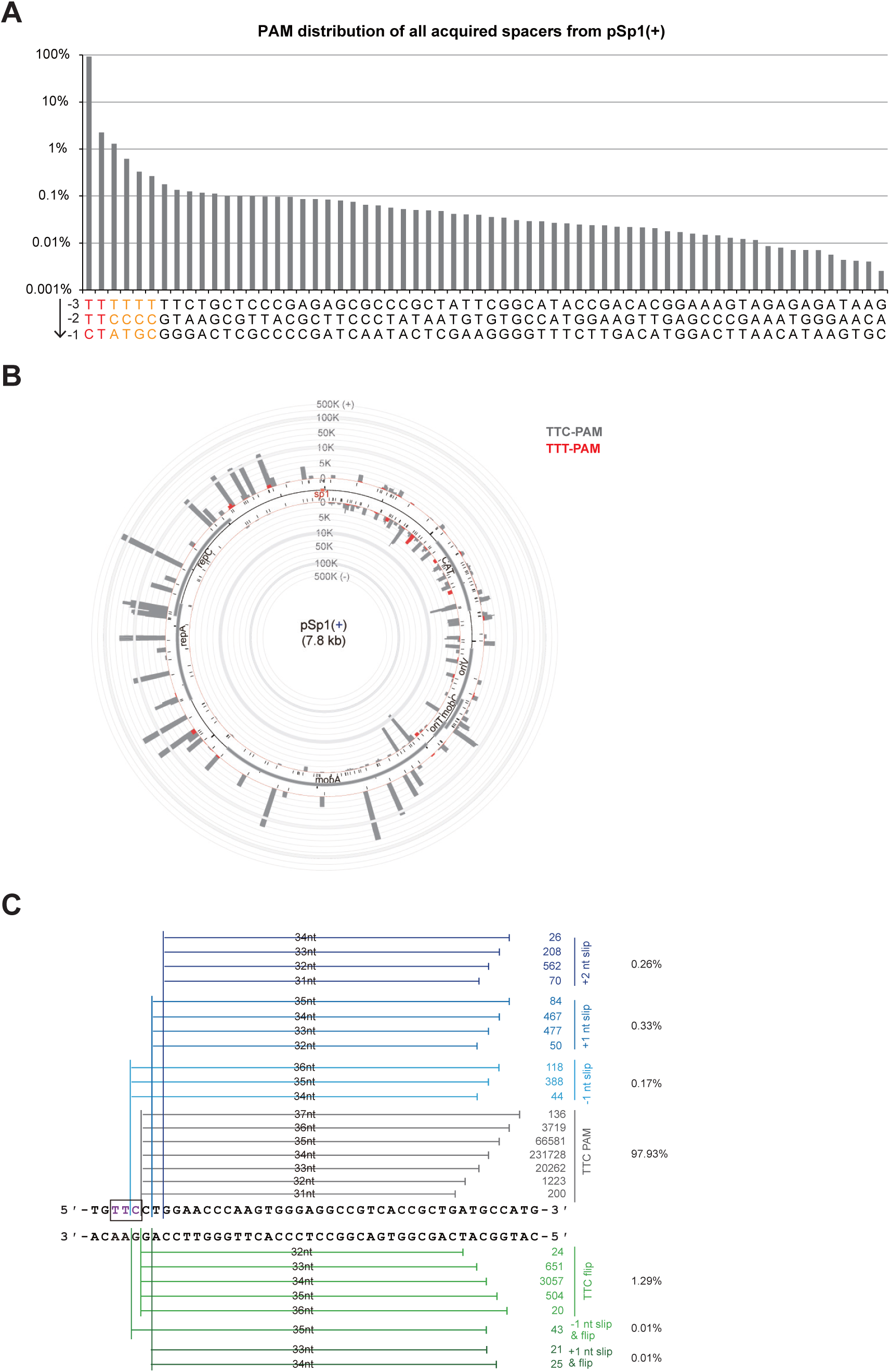
Analyses of acquired spacers from pSp1(+), related to Fig. 3. **A.** PAM frequencies of all acquired spacers derived from pSp1(+). Note that the canonical TTC and alternative TTT PAMs are the two most frequent motifs, followed by TCN motifs that are likely due to +1 nt slips. **B.** Spacers with the alternative TTT PAM (red) showed independent localizations relative to those with the canonical TTC PAM (grey). Note that the plot scale is not continuous (disrupted by grey rings) in order to fully represent a wide range of spacer acquisition efficiencies. **C.** The major spacer acquisition hotspot exemplifies imprecise size selection, slipping (blue) and flipping (green) events. Unique spacers mapped to either strand of the region were categorized and counted regarding their frequencies.

**Supplemental Figure 2:**
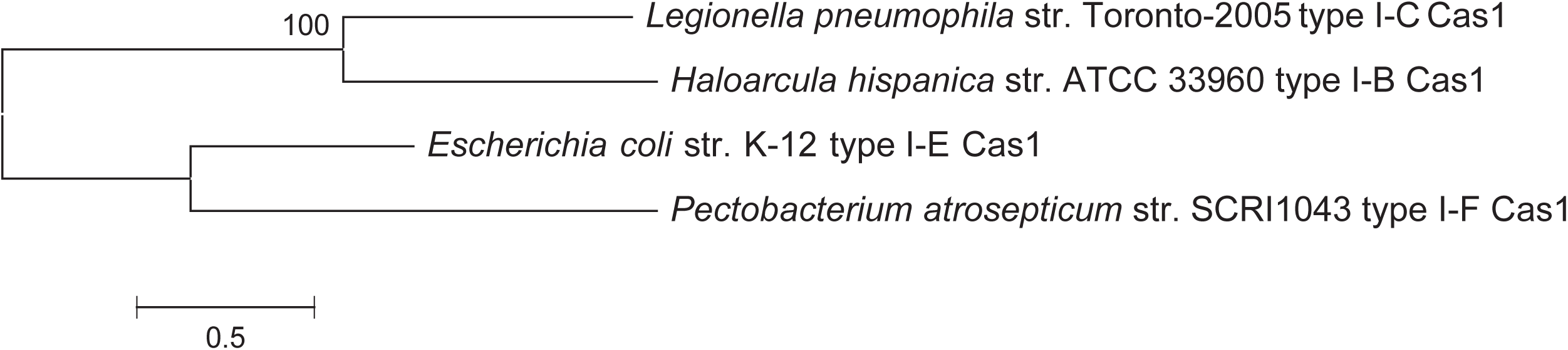
Cas1-based phylogenies of select type I systems. Cas1 protein sequences were retrieved from genomes of *Legionella pneumophila* str. Toronto-2005 (Genbank CP012019), *Haloarcula hispanica* str. ATCC 33960 (Genbank CP002922), *Escherichia coli* str. K-12 (Genbank NC_000913) and *Pectobacterium atrosepticum* str. SCRI1043 (Genbank BX950851). These sequences were aligned using the ClustalW option and subjected to the Maximum Likelihood phylogenetic tree construction using the LG model with 500 bootstrap iterations in MEGA v6.0(Tamura, Stecher et al. 2013).

**Table S1:**
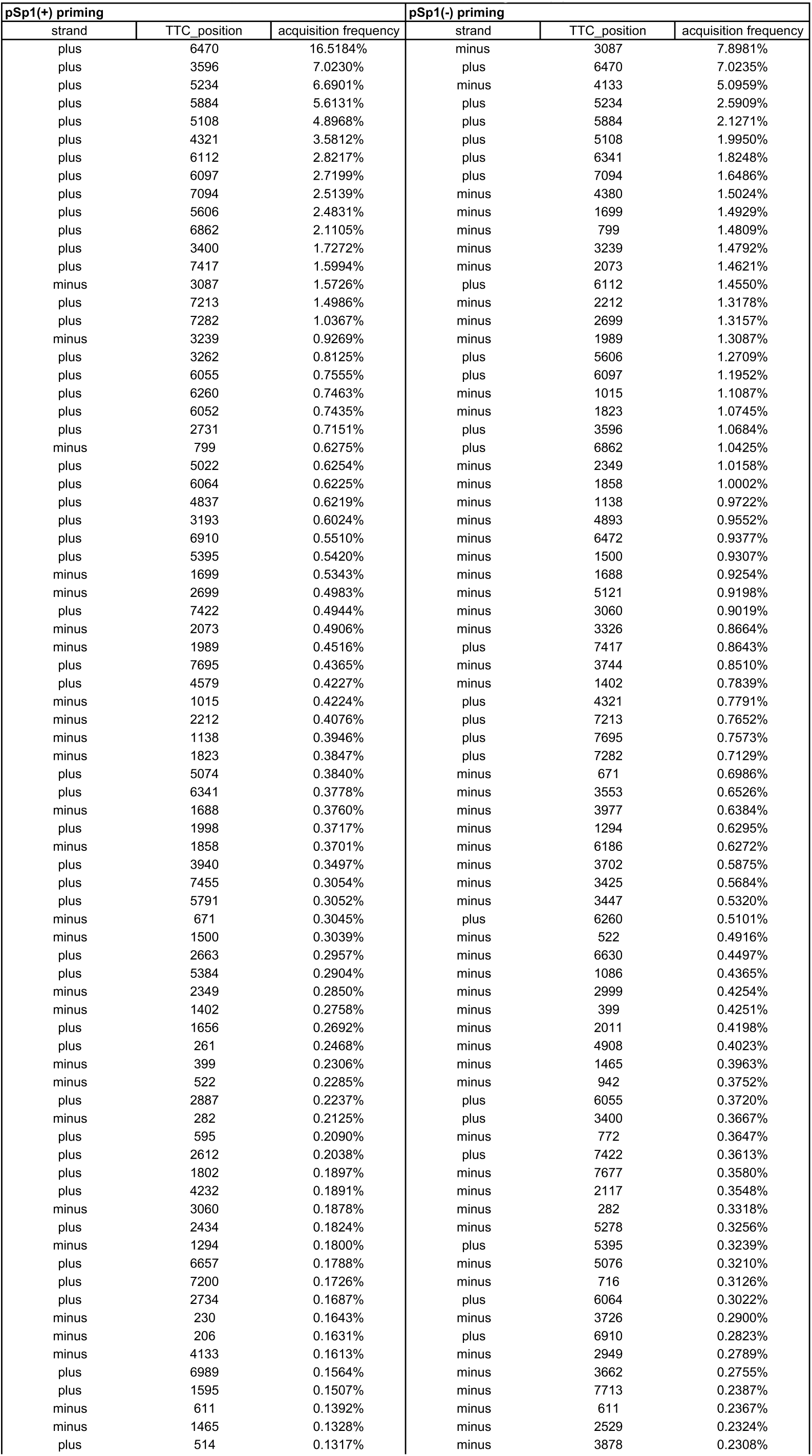
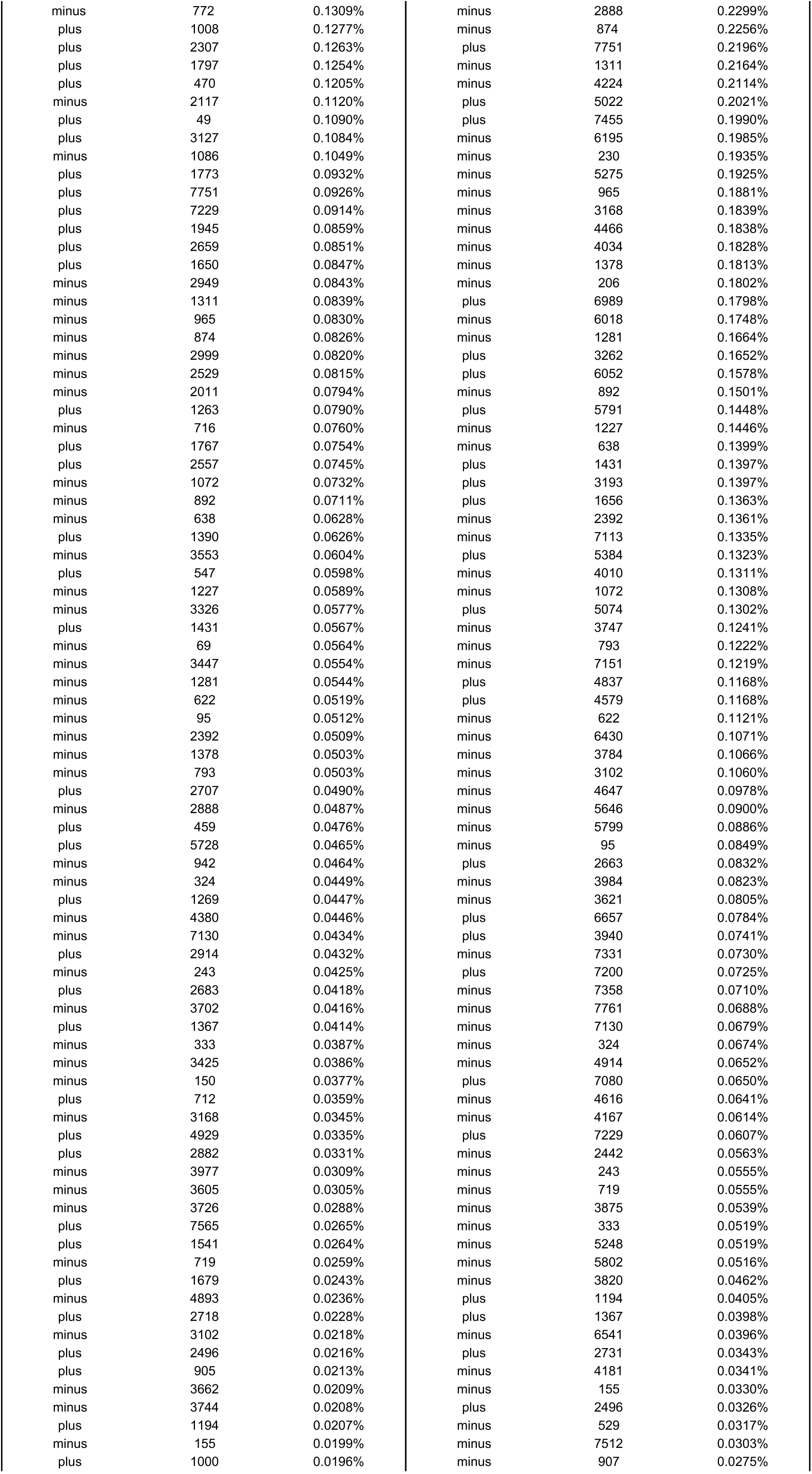
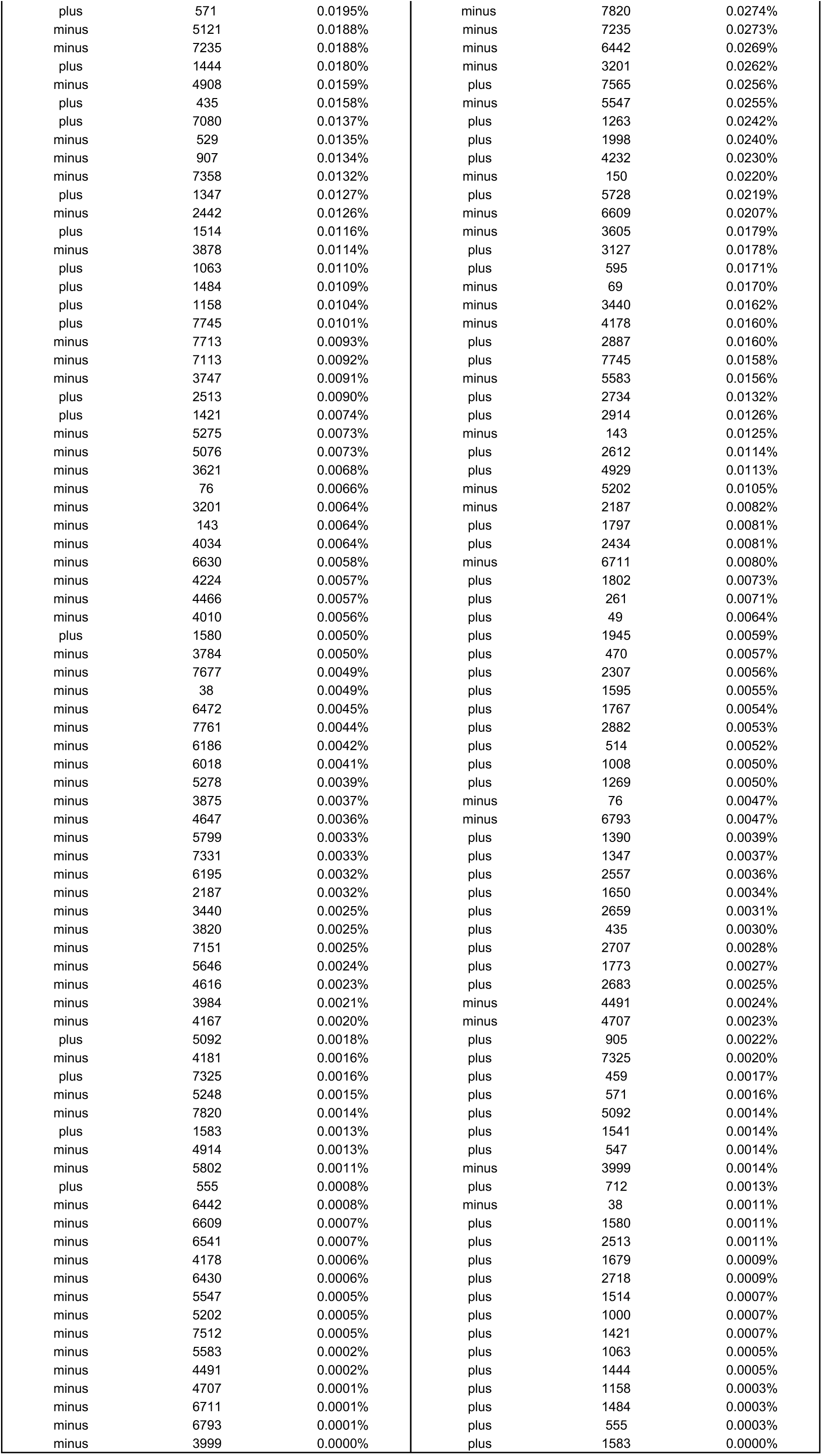
Spacer acquisition frequencies at TTC PAM sites on the pSp1 priming plasmid.

**Table S2:**
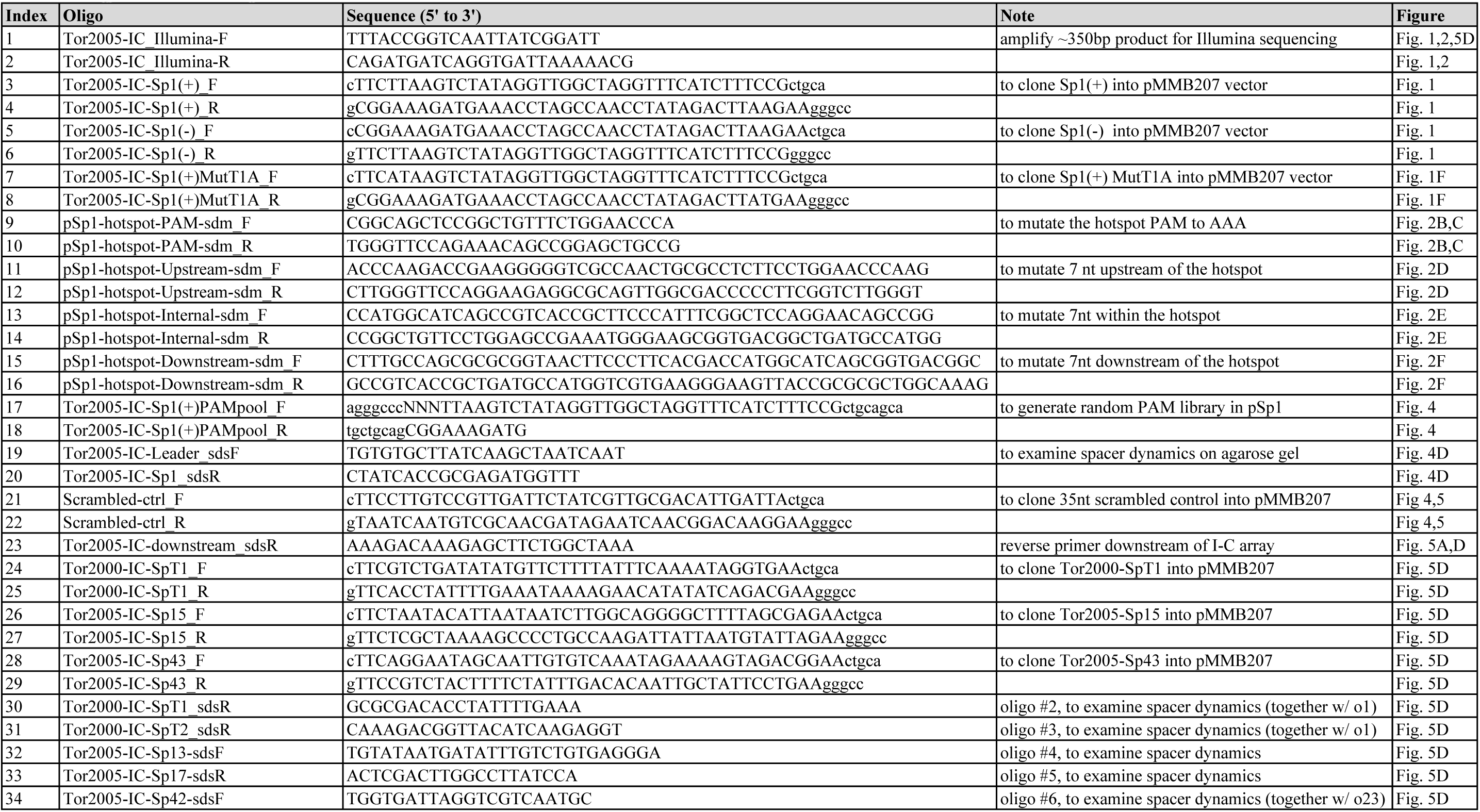
Oligos used in this study.

**Supplemental File:**
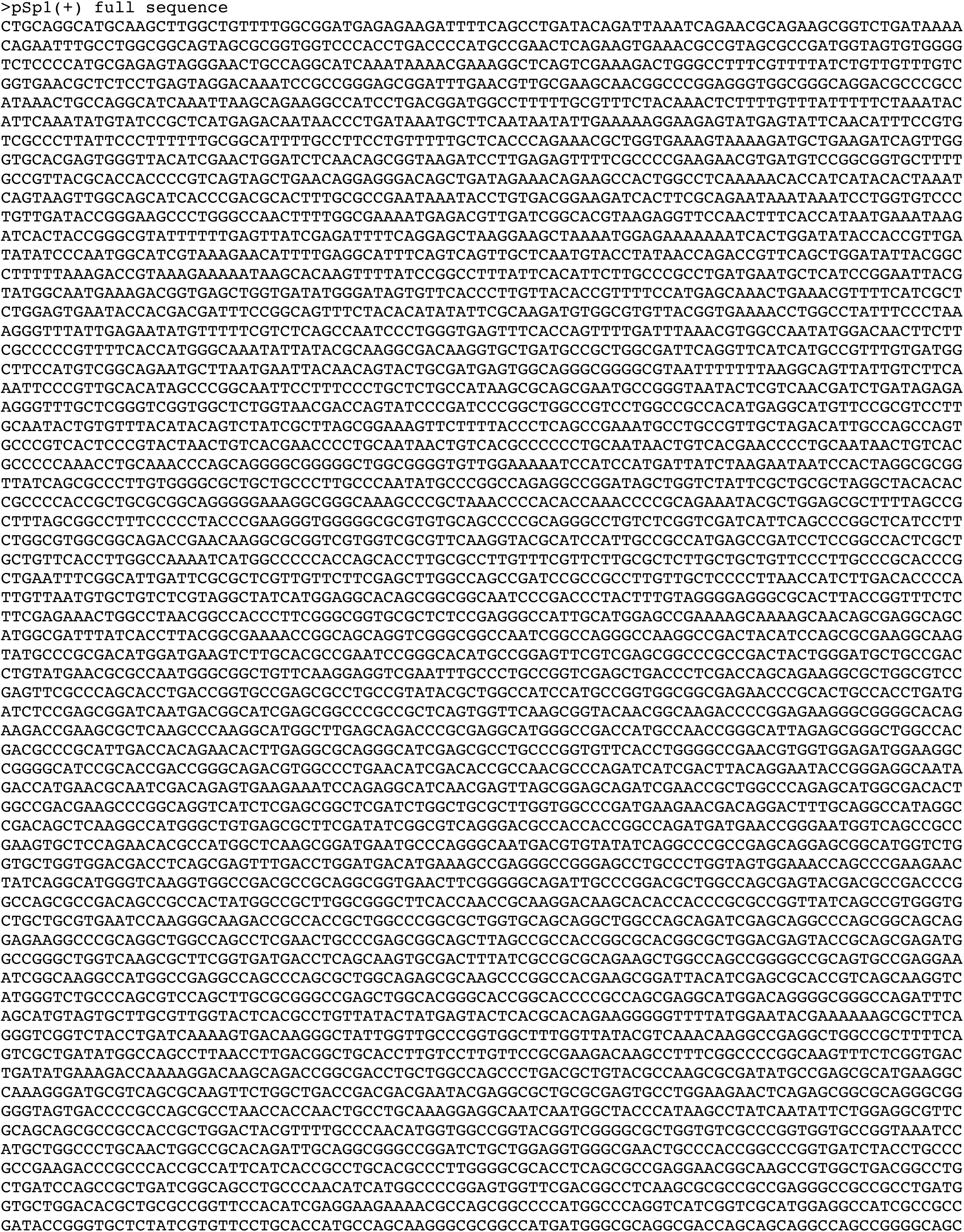
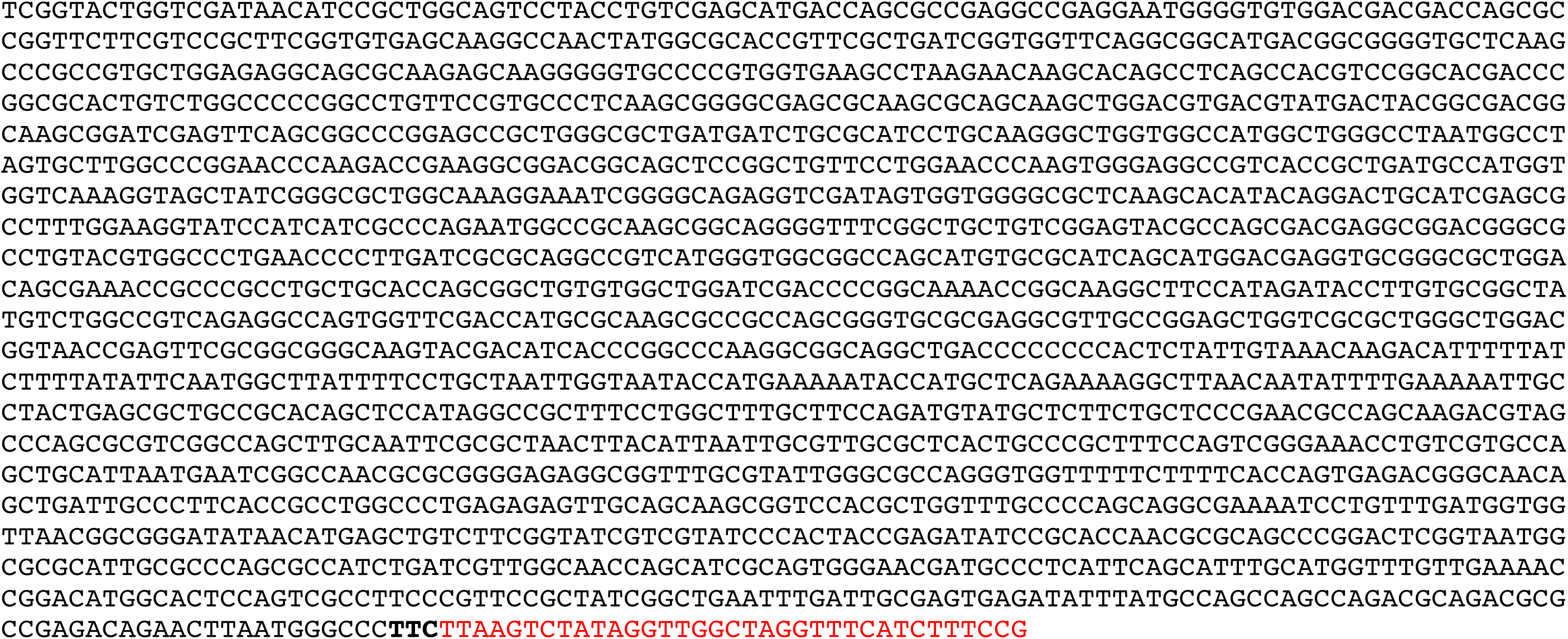
Full nucleotide sequence of pSp1(+).

